# iPSC-derived healthy human astrocytes selectively load miRNAs targeting neuronal genes into extracellular vesicles

**DOI:** 10.1101/2023.04.29.538761

**Authors:** Sara Gordillo-Sampedro, Lina Antounians, Wei Wei, Marat Mufteev, Bas Lendemeijer, Steven A. Kushner, Femke M.S. de Vrij, Augusto Zani, James Ellis

**Affiliations:** Program in Developmental and Stem Cell Biology, Hospital for Sick Children, Toronto, ON, Canada; Department of Molecular Genetics, University of Toronto, Toronto, ON, Canada; Division of General and Thoracic Surgery, Hospital for Sick Children, Toronto, ON, Canada; Department of Psychiatry, Erasmus University Medical Center, Rotterdam, the Netherlands; Department of Psychiatry, Columbia University Medical Center, New York, NY, USA; Department of Surgery, University of Toronto, Toronto, ON, Canada

**Keywords:** Astrocytes, induced Pluripotent Stem Cells, microRNA, extracellular vesicles, RNA binding proteins, Rett syndrome, blood biomarker

## Abstract

Astrocytes are in constant communication with neurons during the establishment and maturation of functional networks in the developing brain. Astrocytes release extracellular vesicles (EVs) containing microRNA (miRNA) cargo that regulates transcript stability in recipient cells. Astrocyte released factors are thought to be involved in neurodevelopmental disorders. Healthy astrocytes partially rescue Rett Syndrome (RTT) neuron function. EVs isolated from stem cell progeny also correct aspects of RTT. EVs cross the blood-brain barrier (BBB) and their cargo is found in peripheral blood which may allow non-invasive detection of EV cargo as biomarkers produced by healthy astrocytes. Here we characterize miRNA cargo and sequence motifs in healthy human astrocyte derived EVs (ADEVs). First, human induced Pluripotent Stem Cells (iPSC) were differentiated into Neural Progenitor Cells (NPCs) and subsequently into astrocytes using a rapid differentiation protocol. iPSC derived astrocytes expressed specific markers, displayed intracellular calcium transients and secreted EVs. miRNAs were identified by RNA-Seq on astrocytes and ADEVs and target gene pathway analysis detected brain related terms. The miRNA profile was consistent with astrocyte identity, and included approximately 80 miRNAs found in astrocytes that were relatively depleted in ADEVs suggestive of passive loading. About 120 miRNAs were relatively enriched in ADEVs and motif analysis discovered binding sites for RNA binding proteins FUS, SRSF7 and CELF5. miRNA-483-5p was the most significantly enriched in ADEVs. This miRNA regulates MECP2 expression in neurons and has been found differentially expressed in blood samples from RTT patients. Our results identify potential miRNA biomarkers selectively sorted into ADEVs and implicate RNA binding protein sequence dependent mechanisms for miRNA cargo loading.

## Introduction

Astrocytes participate in synapse formation and neuron maturation during development and maintain brain homeostasis during adulthood. They are part of the blood-brain-barrier (BBB) which confers on the central nervous system a selective permeability that relies on transcellular mechanisms to transport molecules^1–3^. A key astrocyte feature is their ability to act as communication hubs by establishing astrocyte-neuron and astrocyte-astrocyte networks. These networks act via physical connections between astrocytic neuroligins and neuronal neurexins, through gap junctions between astrocytes, or by secreted factors that regulate synapse formation and maturation^4–6^. Astrocyte released factors are implicated in neurodevelopmental disorders such as Rett Syndrome (RTT), where mutations in the transcriptional regulator methyl-CpG binding protein 2 (MECP2) induce defective neural networks^7,8^. Despite neurons being the most affected cell type due to their high protein levels of MECP2, astrocytes also show an array of phenotypes^9,10^. RTT astrocytes induce non-cell autonomous defects in neuron morphology and firing, thought to be driven by astrocyte released factors ^11–13^.

Astrocytes and neurons also communicate by secreting and internalizing extracellular vesicles (EVs). EVs are double membrane-enclosed vesicles generated in the cytoplasm or the multivesicular body (MVB)/ endosomal compartments, which contain cargo in the form of RNA, proteins and lipids^14^. EVs can regulate transcription and translation upon entering their target cells^15^. They are able to cross the BBB and have been detected in all biofluids using non-invasive methods relying on EV sedimentation by ultracentrifugation, precipitation using commercial kits, or size exclusion chromatography. In addition, EVs can be recovered from conditioned media *in vitro*^16–18^. This feature has propelled the study of EV cargo for its potential as a diagnostic and prognostic tool to define RNA and proteins that are differentially expressed in disease^19,20^. In the context of RTT, EVs isolated from mixed neuronal cultures can rescue RTT neuron networks, and their protein cargo was described as containing signaling proteins that are lacking in the absence of MECP2^21^. In addition, Urine derived Stem Cells were shown to produce EVs with a miR-21-5p cargo that rescued neurogenesis and behaviour in a RTT mouse model^22^. The potential for human astrocyte-derived EV (ADEV) cargo as a prognostic biomarker or as a therapy for neurodevelopmental disorders is unknown.

Sources of human astrocytes to study their ADEV cargo profile are limited. Primary astrocytes are isolated from human cadavers and astrocytes can be derived in vitro from human induced Pluripotent Stem Cells (iPSC). During development astrocytes emerge as a heterogeneous cell population arising from radial glia after neurogenesis is underway at week 24 post-conception in humans^23^. Generating astrocytes from iPSC has proven to be a similarly time-consuming endeavor, often requiring specialized equipment to generate a high yield of functionally mature astrocytic cultures^24^. Recently, a novel differentiation approach has been shown to induce rapid astrocyte differentiation. After generating neural progenitor cells (NPC) and enriching them by cell sorting, stimulation of the astrogliogenic JAK-STAT pathway with BMP4 and LIF generates functional human astrocyte cultures in just 28 days that support neuronal network activity^25^. These human astrocyte cell cultures can be used to isolate EVs and characterize their cargo.

ADEVs contain micro-RNA (miRNA) cargo that is modified in response to cytokines, ATP or presence of pathogenic bacteria, which affect the function of target pathways after internalization by the recipient cell^26–28^. *In vitro* experiments have shown that ADEVs are endocytosed by neurons, and their cargo influences neuronal transcripts and subsequently neuronal function^20,29^. However, the basal miRNA cargo enrichment in ADEVs has not been studied in depth in healthy human astrocytes. Moreover, the use of techniques such as microarrays or qRT-PCR to date captures the most abundant conserved transcripts but precludes global analysis of less abundant miRNAs.

miRNAs are short non-coding RNA sequences that bind to complementary mRNAs to regulate transcript stability, usually by destabilizing mRNA poly-A tails which leads to transcript degradation^30^. Some miRNAs have cell type specificity during development and contribute to post-transcriptional regulation to maintain the equilibrium between mRNA transcription and degradation^31^. Sorting of miRNA cargo into EVs is controlled in part by target mRNA abundance which determines the dispensability of certain miRNAs in a passive mechanism of miRNA disposal inside EVs^32^. At the same time, EV cargo miRNAs contain specific RNA-binding protein (RBP) sequence motifs. RBPs bind these sites to guide miRNAs in the MVB/endosomal compartment to remain in the cell or be sorted via the EXOMotif or other sequences into EVs as cargo^33^.

Here, we generated astrocytes from human iPSC-derived NPCs using a fast differentiation protocol yielding functional astrocytes that secrete EVs which can be isolated using ultracentrifugation. Analysis of miRNA cargo using RNA-seq showed that ADEVs contain significantly enriched miRNAs that participate in pathways regulating neuronal activity. XStreme motif discovery analysis unveiled the presence of RBP sequence motifs that are enriched on ADEV-miRNAs, supporting a sequence-dependent miRNA loading mechanism in ADEVs. Indeed, the most differentially sorted miR-483-5p cargo contains a previously described EXOmotif and is known to target *MECP2*. MiR-483-5p is the most differentially expressed in peripheral blood of RTT patients, consistent with it being a prognostic biomarker of astrocyte function in this neurodevelopmental disorder^34,35^.

## Results

### FACS enriched human NPCs rapidly differentiate to astrocyte fate

To study ADEVs, we employed a simple and time efficient human iPSC differentiation protocol that used a modified 21 day dual-SMAD inhibition treatment to produce NPCs that were purified and then differentiated over 28 days into astrocytes^25^ (Fig.1A). First, we derived two replicates of NPCs (NPC1 and NPC2) from the healthy female human PGPC-14 iPSC line^36^. Once expanded, low passage (P2-3) NPCs underwent fluorescence activated cell sorting (FACS) for markers CD184+/CD44-/CD271-/CD24+ to generate homogeneous progenitor cultures^37^. NPC enrichment ensured homogeneous progenitor (SOX1+, Nestin+) status in all cells, by removing mesenchymal/neural crest cells (CD44+/CD271+), and cells that undergo spontaneous differentiation into (MAP2+) neurons (Fig.1B). Two replicate astrocyte cultures (iPSC-AS) were generated from sorted NPCs by 28-day treatment with BMP4 and LIF (Fig.1A). iPSC-AS (WT1 and WT2) cultures were characterized by western blot for astrocyte specific markers compared to primary human commercial astrocytes (HCA, ScienCell #1800). ALDH1L1 and GFAP isoforms were detected in HCA and both iPSC-AS replicates as expected, and were not present in iPSC, NPC or neuron protein extracts (Fig.1C and Supp.Fig.1A). β-Tubulin was observed in the neuron extracts, with minor β-Tubulin contamination in the HCA and iPSC-AS2 lanes. Immunofluorescence confirmed the expected astrocyte morphology and expression of specific astrocyte markers S100β, GFAP, ALDH1L1 and CD44^38,39^ whereas neuronal markers β-Tubulin and progenitor marker SOX1 were not expressed (Fig.1D).

**Figure 1.**
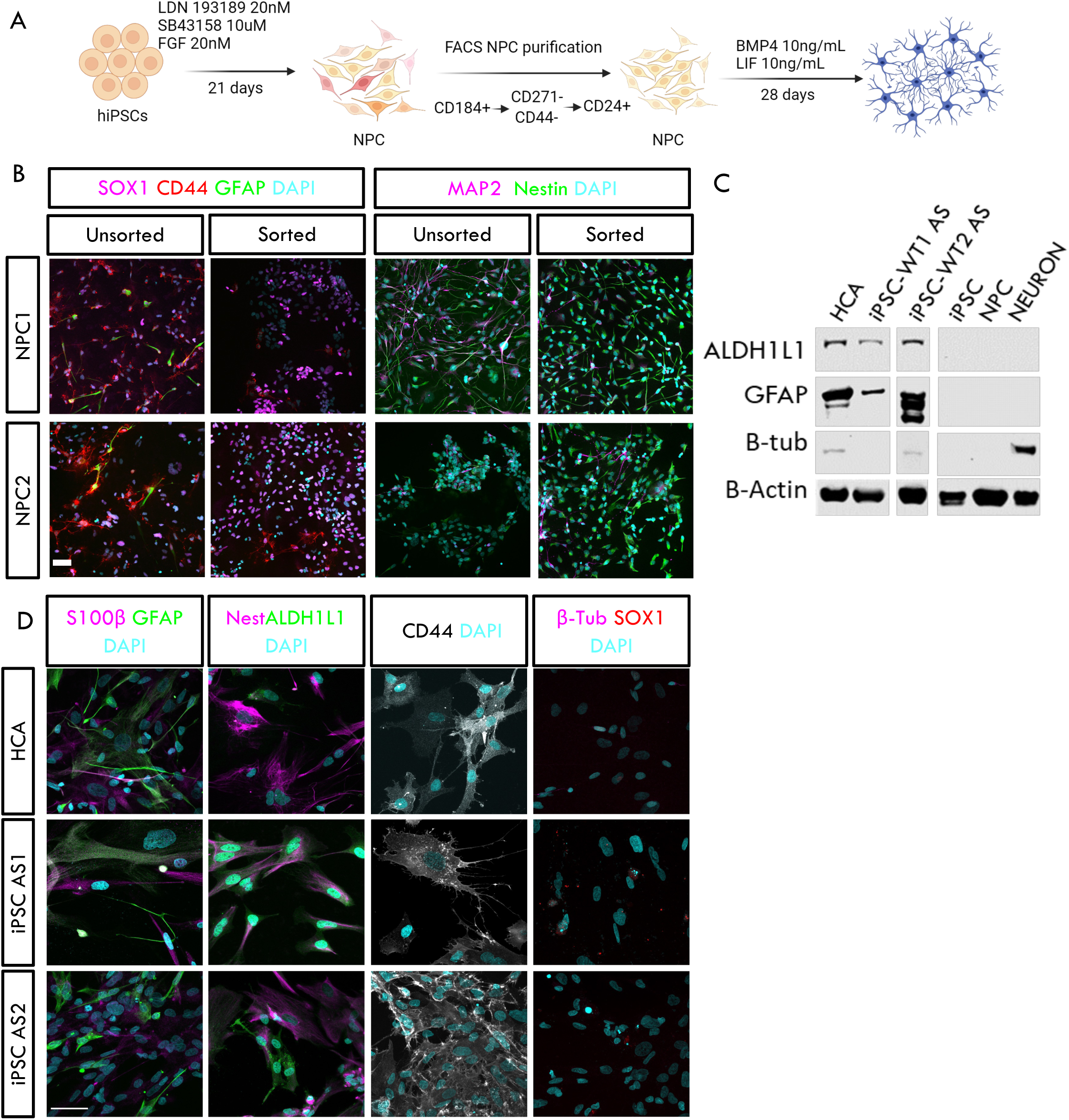
Sorted NPC differentiate into cells expressing astrocyte markers. **A.** Overview of the rapid human iPSC-NPC-astrocyte differentiation protocol. **B.** Cell sorting removes most CD44+/GFAP-populations as well as CD44+/GFAP+ and MAP2+ cells. Some NPCs after sorting express GFAP as well as cytoplasmic MAP2 as previously reported. Images taken at 20x. **C.** Western blot showing iPSC-AS express astrocyte specific markers that are not present in iPSC, NPC or neuron preparations. **D.** iPSC-AS display similar morphology and astrocyte marker expression as HCA. Images taken at 40x. Scale bars 50μm.

### iPSC-AS display characteristic astrocyte activity

To assess iPSC-AS function in culture, we examined intracellular calcium transients as a functional feature of astrocyte networks that generate and propagate intracellular calcium waves spontaneously and in response to stimuli. Calcium transients can be observed in live cultures by incubating cells for 15 minutes in 2μM fluorescence indicator Fluo4-AM^40^ (Fig.2A and Supp.Fig.1B). Baseline recordings of spontaneous calcium transients over a period of 5 minutes showed both HCA and iPSC-AS cultures exhibited intracellular calcium dynamics indicating these cultures display previously reported functional features (Fig.2B, and Supp.Fig1.C). Taken together, these results indicate successful differentiation of iPSC to functional astrocytes that are similar to primary human astrocytes.

**Figure 2.**
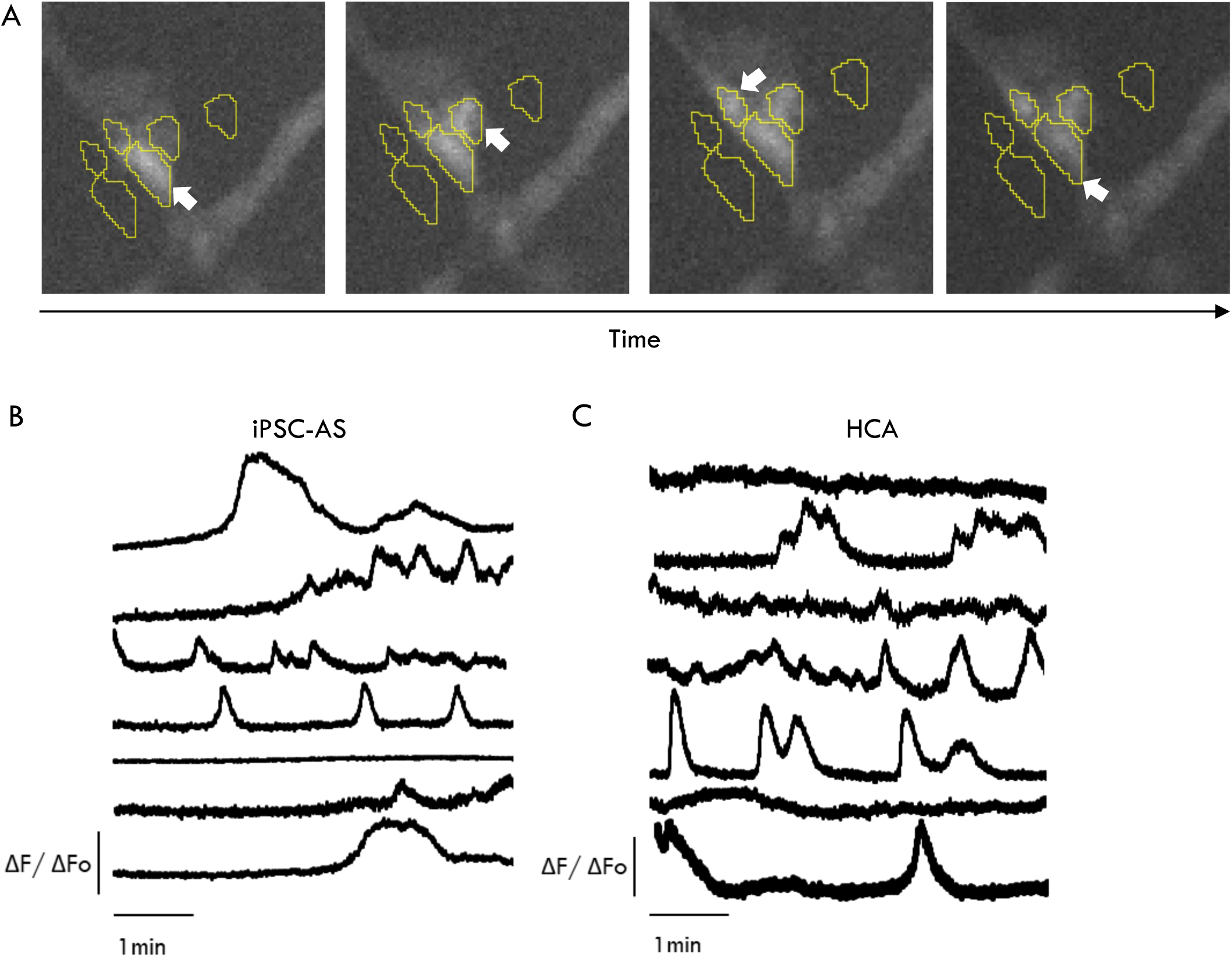
Intracellular calcium transient activity in functional astrocyte networks. **A.** Intracellular calcium labelled with indicator Fluo4-AM to visualize calcium transient propagation (white arrows) between different astrocytes. **B-C.** Representative ΔF/Fo traces showing spontaneous intracellular Ca2+ transients in iPSC-AS and HCA cultures recorded for 5 minutes.

### iPSC-AS secrete ADEVs isolated by ultracentrifugation

ADEVs contain miRNA cargo protected from nucleases^20^. With the aim of studying ADEV cargo, we harvested astrocyte conditioned medium (using exosome depleted fetal bovine serum) to isolate ADEVs from HCA and iPSC-AS following a standard ultracentrifugation protocol^41^ (Fig.3A). Isolated EV-HCA and EV-iPSC-AS displayed the expected size range (50-250nm) using nanoparticle tracking analysis and morphology observed with transmission electron microscopy (Fig.3B)^42^. The ADEV preparations showed no indications of cellular debris contamination detected by an antibody to Golgi network protein GM130 on western blot (Fig.3C). As expected, GM130 was observed in HCA whole cell extracts but not in previously reported EV-hMSC isolated from human Mesenchymal Stem Cells. These results show that ADEVs can successfully be isolated from HCA and iPSC-AS, and exhibit the expected size, concentration and morphology.

**Figure 3.**
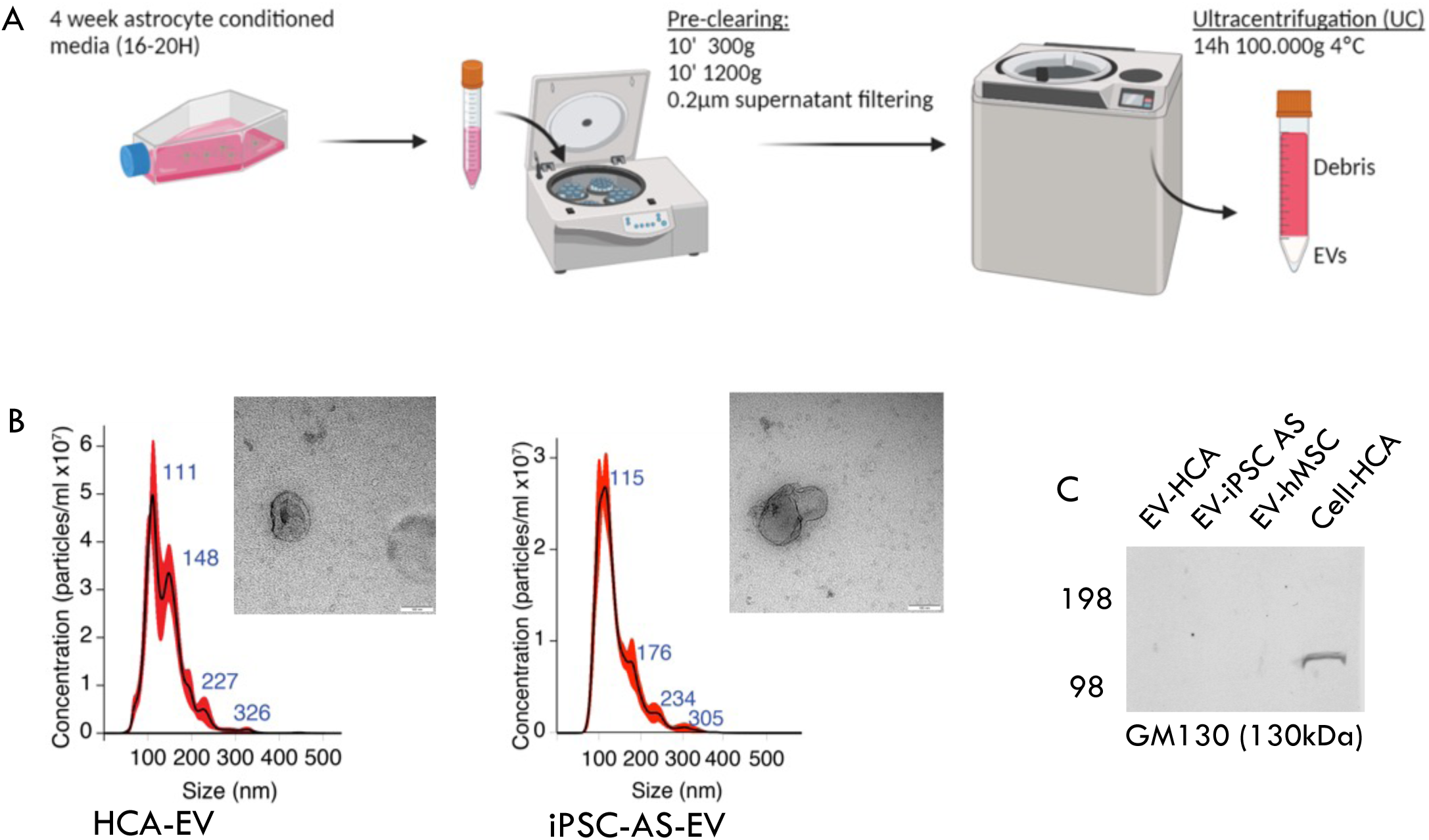
ADEV isolation and quality control. **A.** Overview of ADEV isolation protocol from astrocyte conditioned medium. **B.** NTA quantification and TEM imaging showing HCA-EV and iPSC-AS-EV have expected concentrations over a size range from 50 to 250nm and expected vesicle morphology. **C.** Western blot showing ADEV samples are not contaminated by cell proteins. Golgi network protein GM130 is only detected in the whole cell sample at the expected size of 130kDa.

### iPSC-AS and primary astrocytes express specific miRNAs

With the aim of obtaining a broad picture of miRNA expression in control astrocytes and ADEVs, we used an RNA sequencing strategy. Whole cell RNA was isolated from 2 replicates of HCA and iPSC-AS. For ADEV samples, the small RNA fraction was isolated from 2 replicates of HCA-EV and 3 replicates of iPSC-AS-EV. Samples were sequenced on an Illumina NovaSeq SP flowcell for single end 50bp length reads, aiming for 40M reads per transcript. Raw data was processed using the server based online tool MirDeep2^43^ to obtain miRNA transcript counts for a total of 1709 human miRNAs (Supp.Table1).

Principal component analysis showed that samples clustered based on their origin (Fig.4A), indicating differences between their whole cell and ADEV miRNA profiles (Supp.Fig.2A). First, we compared miRNAs in HCA vs. iPSC-AS to determine how reproducible the signatures are between cultures of different origins. Human primary cell cultures are often heterogeneous and grown in medium that contains fetal bovine serum, a component known to induce reactive phenotypes in astrocyte cultures that affect their transcriptome^44^. Differential gene expression analysis using DESeq2 showed a small subset of miRNAs differentially expressed in HCA vs. iPSC-AS (Fig.4B). To find miRNA expression supporting astrocyte identity we compared miRNAs that were not differentially expressed in HCA vs iPSC-AS so they had similar expression in both cultures. We searched for candidate miRNAs previously reported in the literature, some of which are involved with astrocytic competency acquisition and astrocyte differentiation like miR-153, miR-125 or the let-7 family of miRNAs ^45–47^. The presence of these and other reported astrocyte enriched miRNAs in both HCA and iPSC-AS whole cell samples further support astrocyte identity, albeit with no information regarding their absolute abundance^48^ (Fig.4C). These results show that RNA isolates from HCA and iPSC-AS yield similar miRNA profiles.

**Figure 4.**
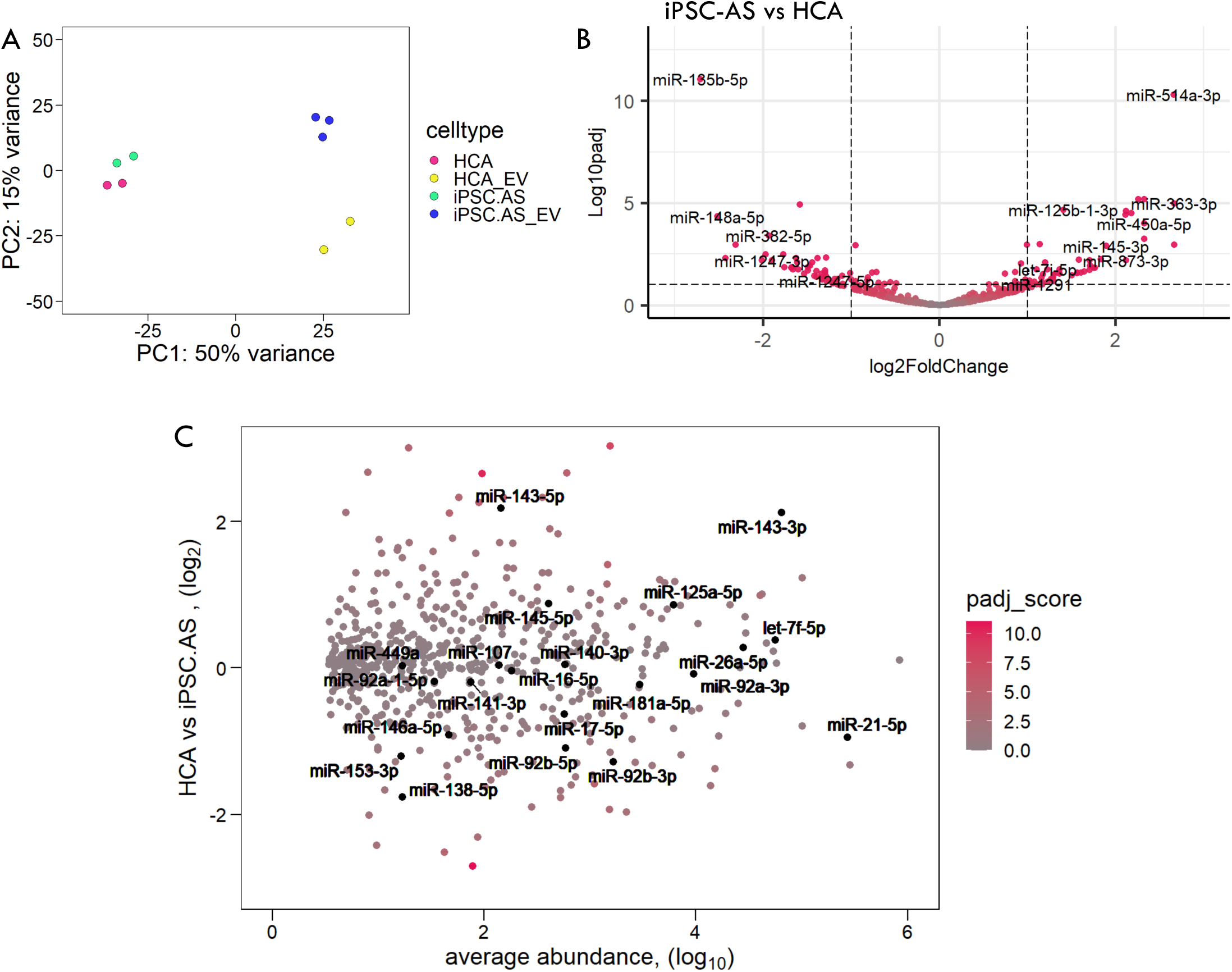
RNAseq analysis on astrocytes and ADEVs. **A.** PCA plot showing variance distribution stems from differences between whole cell astrocytes and ADEV samples. **B.** Volcano plot showing differential miRNA expression of iPSC-AS vs HCA. **C.** Scatter plot of shared astrocyte associated transcripts reported in literature (padj>0.05, grey dots). X-axis represents the average abundance (baseMean) of each miRNA (normalized counts) and Y-axis represents their fold-change in HCA vs iPSC-AS. Color gradient for padj score shows other significant genes in pink. Black dots show miRNAs associated with astrocyte differentiation and identity^45–48^.

### ADEVs selectively sort specific miRNAs that regulate neuronal function

Next, we explored the ADEV miRNA cargo. We first defined a subset of 78 miRNAs depleted in iPSC-AS-EV compared to the whole cell samples (Fig.5A and Supp.Table.2). A similar number of 79 miRNAs were also depleted in the HCA-EV (Supp.Fig.2B and Supp.Table.2). These miRNAs are detected as ADEV cargo, but their relative abundance is very low, suggesting their functional contribution would likely be negligible, excluding them as candidates in transcriptional regulation processes in the target cell. However, these miRNAs could be passively loaded into ADEVs to regulate their abundance in astrocytes, where they could be carrying out important cell-autonomous functions. For instance, elevated levels of let-7 and miR-125 in astrocyte cultures primes progenitor cells for astrogliogenesis by stimulating the JAK-STAT pathway^49^. Thus, ADEV-depleted miRNAs could still yield information on astrocyte phenotypes. On the other hand, we defined a subset of 119 miRNAs enriched in iPSC-AS-EV (padj<0.05)(Fig.5B), including miR-451 and miR-142-3p (Supp.Table.2) shown to be selectively loaded into EVs^50^. Similarly, 155 miRNAs were enriched in HCA-EV (Supp.Fig.2C and Supp.Table.2). Our hypothesis is that EVs are internalized by the surrounding cells including neurons, where their miRNA cargo participates in fine-tuning transcripts in response to environmental cues perceived by astrocytes in their roles as homeostasis regulators. To find indications of potential non-cell autonomous functional roles of ADEV-miRNAs once internalized by their target cells, we ran target gene pathway analysis using the online tool MirSystem^51^ for the top 20 EV-miRNAs. Several brain and developmental terms emerged such as axon guidance and the neurotrophin signalling pathway which regulates cell differentiation and survival as well as synapse formation (Fig.5C). Our data indicate that these miRNAs are likely contributing to neuronal function^26^. In line with our whole cell analysis, both HCA-EV and iPSC-AS-EV showed similar Top 20 enriched and depleted miRNA profiles further supporting replicability between these cultures (Fig.5D).

**Figure 5.**
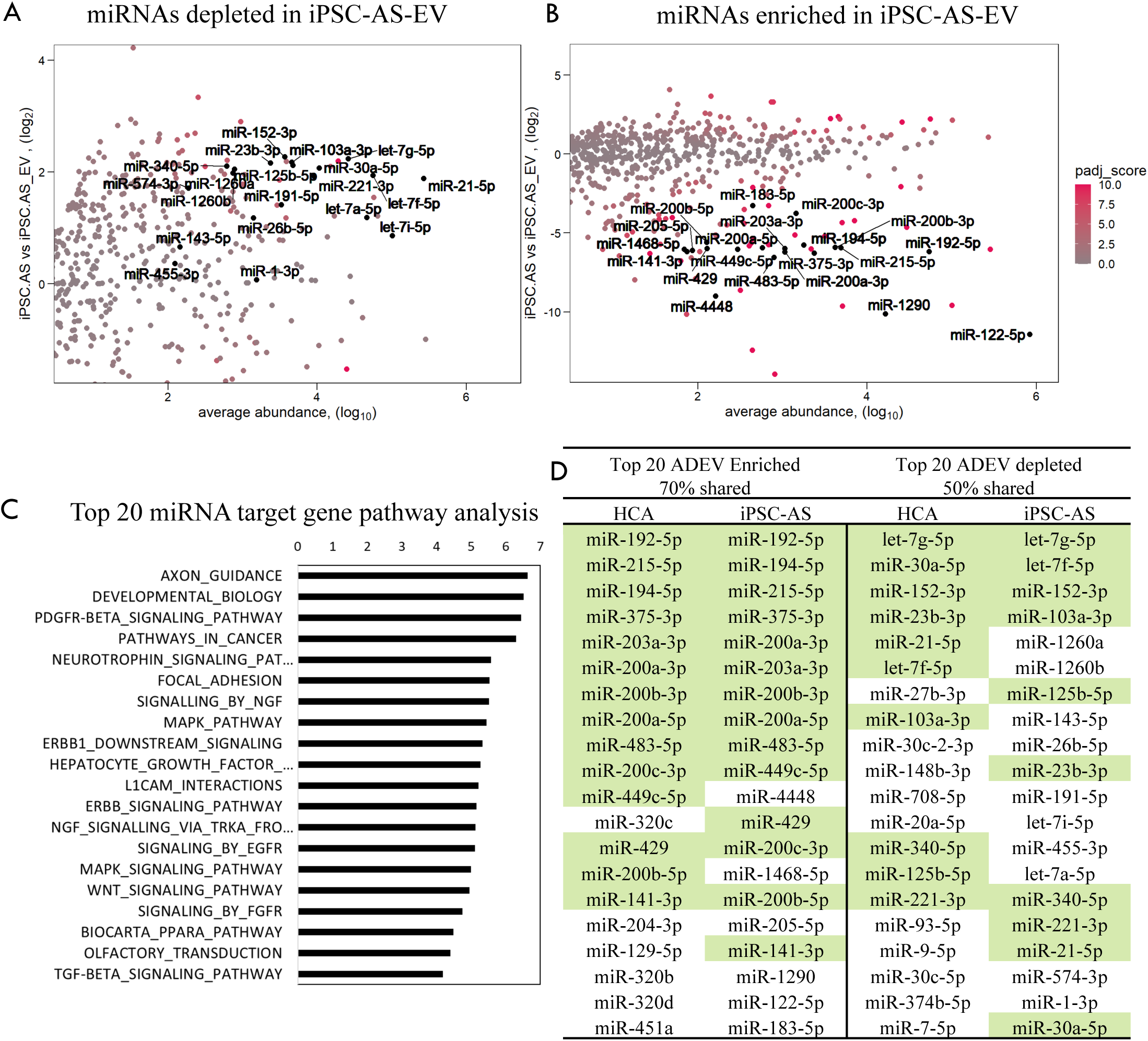
ADEV-miRNA depletion and enrichment analysis. **A.** Scatter plot showing top 20 miRNAs (in black) depleted in iPSC-AS-EV and **B.** Scatter plot showing top 20 miRNAs (in black) enriched in iPSC-AS-EV ordered by p-value. X-axis represents the average abundance (baseMean) of each miRNA (normalized counts) and Y-axis represents fold-change of iPSC-AS/iPSC-AS-EV ratio. Color gradient for padj score (-log(padj, 10)) shows most significant genes in pink. **C.** Target gene pathway analysis of top 20 ADEV enriched miRNAs with relevance score (x axis) assigned based on foldchange values from miRSystem. **D.** List of top 20 enriched miRNAs in HCA-EV (Supp.Fig.2C) and iPSC-AS-EV (Fig.5C), and top 20 depleted miRNAs in HCA-EV (Supp.Fig.2B) and iPSC-AS-EV (Fig.5B) ordered by padj. In green, top 20 enriched or depleted miRNAs found in both HCA-EV and iPSC-AS-EV.

These results show that ADEVs contain a subset of miRNAs that are enriched compared to miRNAs remaining in the astrocytes, suggesting a mechanism of active selection of EV cargo. In addition, the sorted miRNAs target transcripts with relevant functions for correct neuronal maturation, indicating they could be secreted by astrocytes to communicate with surrounding neurons and synapses.

### Enriched miRNAs contain RBP binding motifs for sequence dependent sorting

One of the mechanisms thought to regulate miRNA sorting into EVs relies on RBP sequence dependent interactions. Recently, EXOMotifs have been described in some mouse cell types where RBPs, such as fused in sarcoma (FUS) which participates in miRNA silencing, interact with miRNAs to load them into EVs^33,52^. To find indications of potentially similar mechanisms in iPSC-AS-EV miRNA loading, we analyzed RBP motif enrichment using XStreme motif discovery tool^53^ for sequences between 4-8 nucleotides in miRNAs (Supp.Table.3). We also analyzed motifs in precursor miRNA sequences (pre-miRNAs) which are processed in the same compartment where EVs are loaded^54^. We found enriched motifs for the following RBPs in ADEV-miRNAs (Fig.6A): Serine/Arginine rich splicing factor family 7 (SRSF7); CUGBP Elav-Like Family Member 5 (CELF5/BRUNOL5); FUS; and Sterile alpha motif domain containing A4 (SAMD4A). These RBPs are expressed in human fetal astrocytes, with CELF5 expressed both in neurons and astrocytes^55^. They have reported roles in neuron RNA splicing and export and/or miRNA binding ability, in line with an RBP sequence-dependent loading mechanism and regulation of neuronal transcripts via EV cargo^56–58^. While the small RNA size selection prior to sequencing does not allow detection of pre-miRNAs in ADEVs, we searched sequences of pre-miRNAs corresponding to detected ADEV enriched miRNAs for RBP motifs. Pre-miRNA enriched motifs (Figure 6A) were found for SRSF7, and RNA binding motif 28 (RBM28) or deleted in azoospermia-associated protein 1 (DAZAP1). These RBPs may have some role in pre-miRNA localization or processing in astrocytes. Overall, our computational findings support the model where miRNAs targeting neuronal genes would be selectively loaded into ADEVs via RBP sequence motif recognition.

**Figure 6.**
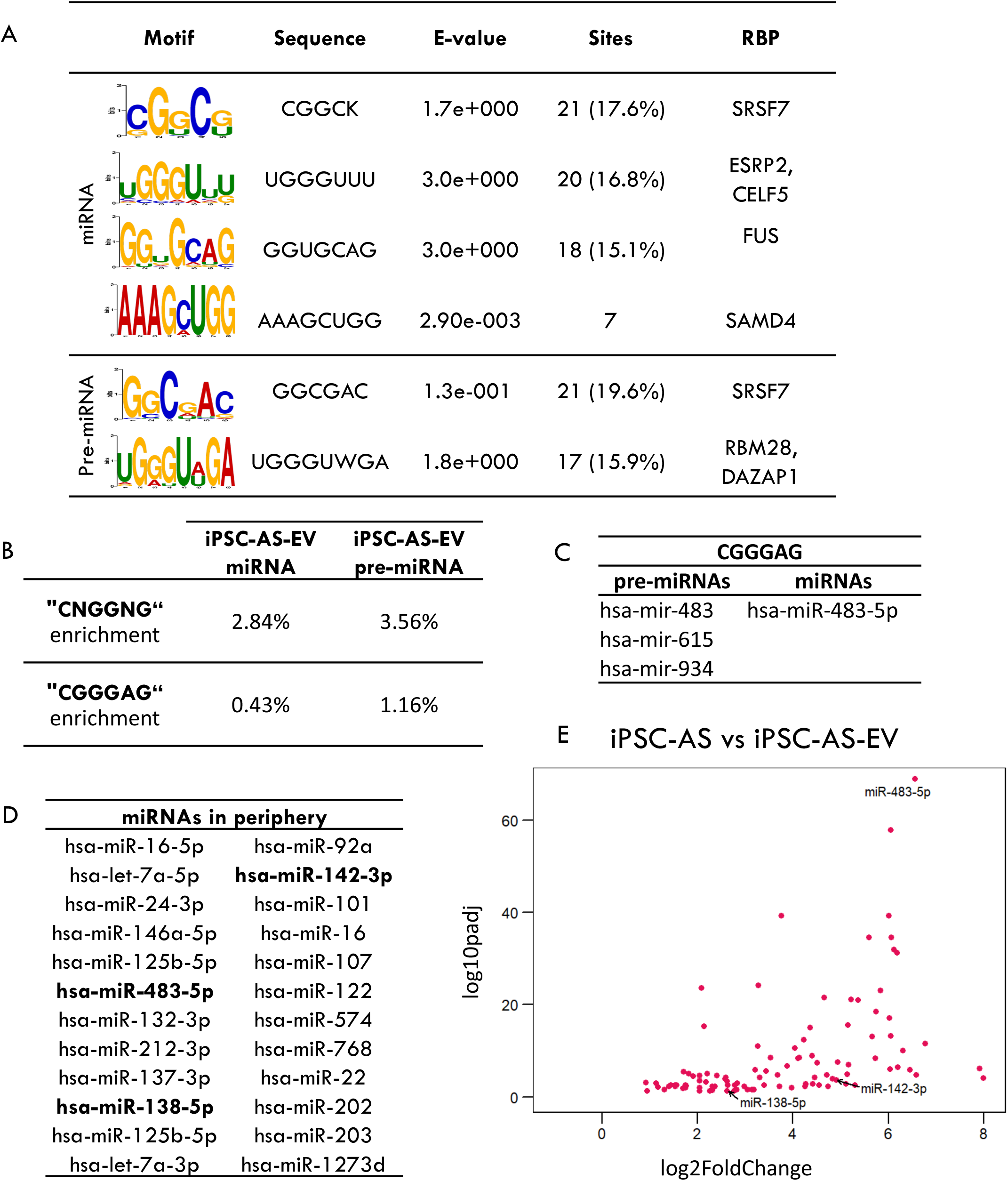
RBP motif discovery on enriched ADEV miRNAs and pre-miRNAs. **A.** XStreme RBP motif discovery results found in mature (miRNA) and precursor (pre-miRNA) sequences. E-value: accurate estimate of statistical significance taking into account the p-value and the number of motifs reported by STREME using 0.05 as threshold. Sites: Number of sequences with the motif (percentage). XStreme provides any RBP known to contain any of the enriched motifs. **B.** Enrichment percentage of miRNAs and pre-miRNAs containing non-cell-type specific EXOMotif reported in *García-Martin et al Nature 2022.* % enrichment = miRNAs with motif/total number of miRNAs detected in ADEVs x100. **C.** Pre-miRNAs and miRNAs containing EXOMotif CGGGAG in iPSC-AS. **D.** List of miRNAs reported in literature of healthy individuals and RTT patients. Bold miRNAs are also shown in E. **E.** Volcano plot showing fold-change enrichment and padj scores of iPSC-AS-EV miRNAs reported in human peripheral samples.

However, our XSTREME based motif discovery did not identify the recently reported EXOMotifs CNGGNG and CGGGAG that are bound by ALYREF and FUS and shown to be sufficient to induce miRNA loading into EVs^33^. In our analysis, the ADEV enriched FUS motif was GGUGCAG which overlaps with the GGNG sequence in the first EXOMotif. To assess the enrichment of these EXOMotifs in EV-iPSC-AS miRNAs, we manually searched for them in miRNA and pre-miRNA sequences. We found little difference in the frequency of enriched ADEV-miRNAs compared with all ADEV-miRNA sequences. The motif CNGGNG showed enrichment of 2.84% and 3.56% in miRNA and pre-miRNA sequences respectively versus the reported 13% enrichment in other cell types (Fig.6B). The CGGGAG sequence was even less enriched. These results are in line with the motifs not being identified in our XSTREME discovery. Differences in enrichment percentage could be due to species or cell type specificity or the use of different methods for motif discovery. Interestingly, miR-483-5p was the only miRNA displaying EXOMotif CGGGAG, suggesting this miRNA is actively loaded into astrocyte EVs via this RBP-sequence dependent mechanism, while 3 pre-miRNAs including mir-483 retained this sequence (Fig.6C). Overall, the EXOMotifs are not the most abundant RBP binding sequences found on ADEV-miRNAs.

### miR-483-5p loaded into ADEVs is a candidate biomarker produced by healthy astrocytes

One of the most exploitable attributes of EVs is their ability to pass through the BBB, which makes them and their cargo interesting candidates in the context of prognostics and therapeutics. To find indications of brain miRNA presence in peripheral samples, we compiled a list of miRNAs reported in blood and saliva samples from healthy individuals^59^ and RTT patients^35^ (Fig.6D). Three ADEV-enriched miRNAs were a match (miR-483-5p, miR-138-5p and miR-142-3p) suggesting brain miRNAs could be recovered in peripheral biofluids. The most significantly enriched of these miRNAs in ADEVs is miR-483-5p (Fig.6E). MiR-483-5p regulates *MECP2* expression in fetal brains^34^ and was found to be downregulated in RTT patient plasma samples^35^. The fact that miR-483-5p contains the EXOmotif CGGGAG suggests this miRNA in particular is actively loaded into ADEVs, perhaps making it to the periphery where it could provide prognostic biomarker information on disorders such as RTT.

## Discussion

We used RNASeq to identify the miRNA profile in healthy human astrocytes, to define the ADEV cargo, and to discover motifs enriched on the miRNAs that are selectively sorted into ADEVs. Computational analysis of the top ADEV enriched miRNAs revealed target pathway terms related to neuronal function. These results indicate that ADEVs are one mechanism of communication between astrocytes and neurons with potential functional consequences for the target cell.

Our motif analysis supported an RBP sequence-dependent mechanism for miRNA loading into ADEVs. Roughly 15% of sorted miRNAs retain consensus binding sites for FUS, an RBP known to mediate miRNA sorting into EVs^33^ via EXOMotifs. This result suggests FUS recognizes an array of EXOMotifs in miRNAs and provides confidence about the accuracy of our motif analysis. In addition, FUS has been implicated in the biogenesis and processing of miRNAs that participate in the regulation of neuronal outgrowth, differentiation and synaptogenesis, suggesting involvement in neuronal function^52^.

Another 17% of sorted miRNAs contain motifs for SRSF7. SRSF7 is reported to bind to and participate in miRNA biogenesis and processing. FUS and SRSF proteins have been found in the secreted proteome of mouse astrocytes^13^ and in EVs from 14 human cell types, suggesting they are secreted together with miRNAs^60^. Interestingly, FUS and SRSF proteins have been shown to play important roles in the pathogenesis of the neurodegenerative disorder amyotrophic lateral sclerosis (ALS), characterized by progressive loss of motor function with 10% of the cases following an inherited familial form with mutations in genes such as FUS. Relevant to our astrocyte study, mutant FUS and another SRSF protein, SRSF1, are involved in astrocyte mediated neurotoxicity in a model for ALS^61^. In addition, 17% of miRNAs bore enriched motifs for CELF5. The CELF family of proteins participate in RNA splicing and transcript stability. They have been studied in the context of neurological disorders such as Fragile X Syndrome, where deficits in astrocyte released factors disrupt synapse formation and maturation^62^. Overall, our analysis predicts that healthy astrocytes selectively sort miRNAs with motifs for FUS, SRSF7 and CELF5 into ADEVs.

We also analyzed RBP motifs in pre-miRNA sequences. About 16% of transcripts contain motifs for DAZAP1 or RBM28, and 19% for SRSF7. This could indicate the involvement of these RBPs in miRNA processing, shown to happen in the MBV^54^. Pre-miRNAs being loaded into EVs have not been reported to date^18^. Alternatively, any of our reported RBPs could be binding ADEV-miRNAs in the target cell as a mechanism to regulate neuronal function.

The strongest evidence for selective sorting into EVs was found for miRNA-483-5p. This was the only miRNA in ADEVs with the canonical EXOmotif CGGGAG and it was the most significantly enriched in iPSC-AS-EV. We infer that miRNA-483-5p is likely also bound by FUS and its selective sorting into ADEVs facilitates non-cell autonomous communication by astrocytes with neurons. miR-483-5p specifically suppresses human *MECP2* expression in early developmental stages until the early postnatal stage, when synaptic maturation takes place and neuronal MECP2 levels increase^35^. The ADEVs would likely also cross the BBB, to communicate through the blood with other tissues where MECP2 is expressed.

Differences in abundance of miRNAs detected in peripheral samples are associated with several neurological disorders and are thought of as a reflection of disruptions in the brain. Thus, changes in certain miRNAs originating in the brain can be detected in patient biofluids and used for early diagnosis, prognosis and designing therapeutic strategies. miR-483-5p is downregulated in plasma from RTT patients and has been associated with growth delay phenotypes^35^. If astrocytes are the primary source of miR-483-5p in blood, we infer that RTT astrocytes may sense low MECP2 levels and downregulate miRNA-483 expression or its processing. RTT astrocytes may also have reduced sorting of miRNA-483-5p into EVs. Ultimately these effects would communicate to RTT neurons that they should upregulate *MECP2*. Less miRNA-483-5p would cross the BBB leading to reduced detection in the blood as a biomarker that RTT astrocytes are not healthy.

Since miR-483-5p is not exclusively expressed in the brain, with other research showing its involvement in several cancers and adipose tissue^63^, it will be important to test the ADEV cargo of human RTT astrocytes. The pipeline described here to rapidly produce human astrocytes to isolate ADEVs can be used on RTT iPSC-AS to examine changes in their cargo and to investigate their interactions with RBPs that sort miRNAs into ADEVs. This might best be accomplished using RNASeq normalization strategies such as spike-ins to obtain information on absolute miRNA abundance^64^, with a special emphasis on miRNA-483-5p depletion to evaluate its utility as a prognostic biomarker. Such a blood biomarker may be most useful as an early indicator that RTT patient astrocytes are responding in a clinical trial in order to justify continued treatments of IGF1 or other novel therapies. Given reports that EVs can partially rescue RTT neuron function, there may also be value in evaluating the therapeutic potential of ADEVs for RTT or other neurodevelopmental disorders.

## Methods

### NPC purification

Two independent NPC preparations were generated as biological replicates from the human iPSC line PGPC-14 and maintained as previously described^36,65^ in Advanced DMEM/F12, 1% N2, 2% B27 (all Thermofisher Scientific), 1ug/ml Laminin (Sigma-Aldrich), 1% Penicillin-Streptomycin (Gibco) and 20ng/ml basic Fibroblast Growth Factor (Peprotech). For NPC purification, passage 2-3 NPCs were detached with Stempro Accutase cell dissociation reagent (A1110501, Life Technologies) and kept in cold PBS 1%FBS for the duration of the protocol. For single stain and fluorescence minus one (FMO) control samples, between 500.000-2million cells were resuspended in 100ul of PBS 1%FBS for antibody staining. For the sample tube, between 10-40 million cells were resuspended in 200-500ul of PBS 1%FBS for CD184, CD24, CD44, CD271 and Dapi staining. Antibody mixes were incubated for 20-30 mins on ice in the dark and samples were washed with PBS 1%FBS 3x. On the last wash, cells were resuspended in 500ul-700ul of PBS 1%FBS and filtered through 700µm mesh into a 5ml polypropylene tubes (BD352063). Cells were sorted for CD184+/CD44-/CD271-/CD24+^37^ in MoFlo XDP Cell Sorter (Beckman-Coulter) at the SickKids Flow Cytometry facility into Matrigel (Corning) coated 6well plates in a density of 300,000 cells per well, kept in NPC medium^25^ with 20 ng/ml basic Fibroblast Growth Factor added fresh (Peprotech, 100-18c) at 37°C and 5%CO2 until confluent, and then expanded and cryopreserved in liquid nitrogen in 500µl FBS 10%DMSO solution.

### Astrocyte differentiation and culture

Confluent purified NPCs from both biological replicates were passaged using Stempro Accutase cell dissociation reagent (A1110501, Life Technologies), in a 1:4 ratio to new Matrigel (Corning) coated wells in the NPC medium and differentiated to astrocytes as previously reported^25^ by addition of 10ng/ml Bone Morphogenetic Protein 4 (Biovision) and 10ng/ml Leukemia Inhibitory Factor (Peprotech). Cells were passaged an average of once a week for 28 days and medium was changed every other day. Human Commercial Astrocytes (ScienCell #1800) were plated on poly-L-Ornithine (Sigma) coated plates with Astrocyte Medium (ScienCell #1801) and used immediately or passaged one time only using trypsin/EDTA following manufacturer instructions. Medium was changed every other day until 70% confluent and every 3 days after that.

### Immunofluorescence

Cells were plated on 24 well plates (82426, Ibidi) or 19mm glass coverslips coated with Matrigel or Poly-L-Ornithine/laminin and fixed in 4%PFA for 8 minutes when confluent. Afterwards, PFA solution was discarded and cells were washed in PBS 3X and kept in PBS at 4°C until staining. For staining, wells were washed in PBS 3x 5minutes and subsequently incubated with primary antibodies either 1h RT or over night at 4°C in antibody mix solution pH7.4 (for 100ml: 10mL 1x triton, 5mL Tris-HCl pH7.5, 0.9grNaCl, 0.25gr porcine gelatin (G8150, Sigma) and sterile water). After primary incubation, cells were washed 3x 5minutes with PBS and subsequently incubated for 1hRT in PBS with secondary antibodies and Dapi. Finally, cells were washed 3x 5minutes in PBS and kept in PBS at 4°C dark until imaged. Cells were imaged using a Zeiss LSM 880 Airyscan microscope at the PGCRL imaging facility 40x/1.1 (W) immersion and 20x/0.8 objectives as Z-stacks and images were processed using Fiji (ImageJ). See Table1 for antibody details.

### Western blot

Protein was extracted from confluent wells washed in ice-cold PBS using radioimmune precipitation assay (RIPA) buffer (25 mM Tris-HCl, pH 7.6, 150 mM NaCl, 1% Nonidet P-40, 1% sodium deoxycholate and 0.1% SDS), incubated 30 minutes with interspersed vortexing and stored at −80°C until quantified. Protein samples were quantified using the Bio-Rad DC assay kit. Equivalent masses of each protein sample were loaded in individual lanes of a BOLT 4-12% Bis-TRIS gradient gel, and separated at 160V for approximately one hour. Proteins were transferred to nitrocellulose membrane overnight at 60V at 4°C. Following transfer, membranes were rinsed in RO water, then blocked in 5% milk or BSA in PBS for one hour at room temperature shaking. Prior to specific marker staining, total protein was visualized using Revert 700 Total Protein Stain (926-11011, LiCor) and subsequently washed by incubating in Revert wash solution (0.1MNaOH, 30%MeOH in MilliQ water) for 10 minutes. Primary antibody staining was done either 1h RT or over night at 4°C shaking, diluting antibodies in 5% milk or BSA PBS-T. Subsequently, membranes were washed 5 minutes 3x in PBS-T and secondary antibody incubation was carried for 1h at RT shaking. Membranes were washed 3x PBST for 5minutes prior to being imaged using LI-COR Odyssey CLx. See Table1 for antibody details.

### Fluo4-AM live calcium imaging

Astrocytes were plated on Matrigel coated 24 well plates (82426, Ibidi) and cultured until confluent. Confluent wells were washed with PBS and incubated in medium containing Fluo4AM (F14201, Thermo Fisher) with a final concentration of 2µM for 15 minutes at 37°C 5%CO2. Afterwards, cells were washed with PBS and kept in astrocyte medium while imaging. Nikon TE-2000 epifluorescence microscope at the PGCRL imaging facility Solent Scientific enclosure was equilibrated to 37 °C for 30 minutes prior to imaging. Cells were imaged using Hamamatsu Orca-R2 camera and the 20x/0.5 magnification, at the lowest laser intensity for 5 to 8 minutes at maximum speed. Recordings were analyzed using Fiji’s ROI manager tool. Regions of interest were drawn of individual cells and size matched background regions, and fluorescence intensity was measured per time frame. Further processing was performed using Excel. Normalized fluorescence intensity was calculated as the ratio between ROI fluorescence intensity and its corresponding background per time frame. Time frames were transformed to time measures based on the speed of the recording to generate plots of normalized fluorescence intensity over time. Percentage of active cells was calculated manually by counting the number of cells displaying at least one fluorescence intensity oscillation during the recording.

### ADEV isolation and characterization

Astrocyte cultures for ADEV isolation were derived from the two biological replicates of NPCs together with one technical replicate from the second NPC preparation initiated one week later. The astrocytes were washed in PBS and cultured in Advanced DMEM-F12 7.5% EXOsome-depleted FBS (ThermoFisher, Waltham, MA) over night. Astrocyte conditioned media was harvested and residual cell debris was removed by centrifugation at 300 × g followed by 1200 × g both for 10 minutes at room temperature. Supernatant was filtered using 0.20 µm cellulose filter (Corning, NY) and kept frozen at −20 °C until ultracentrifugation. ADEV isolation was performed by ultracentrifugation at 100,000 × g for 14 h at 4 °C (swing bucket rotor on minimum acceleration/break setting, SW 32Ti Beckman Coulter, Brea, CA), and ADEVs were resuspended in 200μL PBS and stored at −20°C until used^16,41^. ADEV characterization was performed using transmission electron microscopy and NanoSight particle tracking analysis as previously described^16,41^, and western blot as described above.

### Whole cell and small RNA isolation for RNAseq

One confluent well of a six well plate from two independent PGPC14WT astrocyte differentiations, or 2 wells of HCA were harvested by scraping cells off in 500µl of PBS and centrifuging them at 1000g for 5 minutes. RNA was isolated from cell pellets and ADEV isolates using SPLIT RNA extraction Kit (LEXOgen) according to the manufacturer’s instructions. For the whole cell extracts instructions for total RNA isolation were followed whereas ADEV sample RNA isolation was enriched for small RNAs, generating two fractions of large RNA and small RNA. RNAs were eluted in 10-15 µl of elution buffer and for ADEV samples, with 1U/µl SUPERaseIn RNase inhibitor (AM2694, Invitrogen) and stored at −80 °C until submitted for sequencing. RNA concentrations were analyzed using Nanodrop for whole cell samples.

### RNA sequencing and data analysis

For whole cell samples, 1µg was provided The Centre for Applied Genomics. For ADEV samples, all isolates were provided for bioanalyzer and sequencing. Sequencing libraries were made using the Small RNA library preparation kit NEBNext (NEB) Small RNA library prep set according to the manufacturer’s instructions. Sequencing was performed using Illumina NovaSeq SP flowcell for single end 50bp length reads, aiming for 40M reads per transcript. Library preparation and sequencing were done at TCAG. Raw sequencing data was processed into count data using the open source web-based platform galaxy.org MiRDeep2 tool as previously, mapping reads to hg38 human genome annotation and mature and precursor miRNA sequences from miRbase database^43,64^. To generate a counts table for differential gene expression, miRNA counts for same mature miRNA from different precursor origin were averaged. Differential expression analysis was performed using DESeq2 package in R. Quality of samples was analyzed using principal component analysis (PCA) in DESeq2 package and pheatmap function was used to display variance between samples and replicates. Enriched miRNAs were obtained by ordering data with padj<0.05 and log2FoldChange<0 for whole cell vs ADEV. ADEV-depleted miRNAs were defined as differentially expressed miRNAs in ADEV samples enriched in whole cell (padj>0.05, log2FoldChange>0 comparing whole cell/EV). Top 20 significant ADEV-enriched miRNAs and their log2FoldChange values were input into miRSystem online tool to generate target gene pathway analysis scores^51^. miRNAs reported in peripheral samples were obtained from literature^35,59^ and the ones also found in our ADEV-miRNAs are displayed in Fig.6D.

### RBP and EXOMotif discovery analysis

ADEV-enriched EXOMotif CGGGAG was obtained from literature^33^. To find this motif and run motif discovery, ADEV-enriched miRNAs were matched to their mature or precursor sequences obtained from miRBase (https://mirbase.org). EXOMotif search was performed in RStudio, and percentage of miRNAs was calculated as the ratio of miRNAs containing the motif vs all detected miRNAs (baseMean>0) as the background. Motif discovery analysis was performed in web-based platform MEME-Suite 5.4.1 using the tool XStreme for motif discovery and enrichment analysis setting width parameters for 4-8 nucleotides^33,53^. All detected pre- or miRNAs (baseMean>0) were used as background. RBP motifs were searched in CSIBP-RNA Single Species RNA (Homo Sapiens).

**Table1.** Antibody list.

| *Antibody* | *Concentration* | *Company and Catalog number* |
| --- | --- | --- |
| *CD184-APC* | 1/250 | BD bioscience cat# 560936 |
| *CD44-FITC* | 1/100 | BD bioscience cat# 560977 |
| *CD271-PE* | 1/250 | BD bioscience cat# 560927 |
| *CD24-PECy7* | 1/250 | BD bioscience cat# 561646 |
| *Dapi* | 1/300 | Sigma Aldrich D9542 |
| *GFAP* | 1/1000 (ICC)<br>1/2000 (WB) | Millipore AB5804 |
| *CD44* | 1/500 | BD Pharmigen cat#550538 |
| *S100β* | 1/1000 | Sigma S2532 |
| *ALDH1L1* | 1/200 (ICC)<br>1:5000 (WB) | Abcam ab190298 |
| *Nestin* | 1:500 (ICC)<br>1:1000 (WB) | Millipore AB5326 |
| *SOX1* | 1/200 | R&D AF3369 |
| *TujIII* | 1/2000 (WB) | Millipore AB1637 |
| *MAP2* | 1/1000 | SYSY 188-004 |
| *βActin* | 1/4000 (WB) | Sigma A5441 |
| *GM130* | 1:500 (WB) | BD Biosciences 610822 |
| *Donkey anti-rabbit Alexa488* | 1/500 | Thermo Fisher A21206 |
| *Donkey anti-rat IgG (H+L)* | 1/500 | Sigma SA5-10027 |
| *Donkey anti-mouse Alexa 647* | 1/500 | Jackson Immuno.715-605-151 |
| *Donkey anti-mouse Alexa 488* | 1/500 | Jackson Immuno. 715-545-150 |
| *Donkey anti-goat Alexa 555* | 1/500 | Thermo Fisher A-21432 |
| *Donkey anti-guinea pig Alexa 647* | 1/500 | Jackson Immuno.706-605-148 |
| *Goat anti-mouse IRDye 800CW* | 1/20.000 | LiCor 926-32210 |
| *Goat anti-rabbit IRDye 800CW* | 1/20.000 | LiCor 925-32211 |

## List of Abbreviations

AS: Astrocytes
ADEVs: Astrocyte Derived Extracellular Vesicles
BBB: Blood Brain Barrier
EV: Extracellular Vesicles
FACS: Fluorescence Activated Cell Sorting
HCA: Human Commercial Astrocytes
iPSC: Induced Pluripotent Stem Cells
miRNA: MicroRNA
MVB: Multivesicular Body
NPC: Neural Progenitor Cells
RBP: RNA Binding Proteins
RTT: Rett syndrome

## Declarations

## Ethics approval and consent to participate

The PGPC-14 iPSC line was previously generated, and was cultured for this study, under the approval of the SickKids Research Ethics Board and the Canadian Institutes of Health Research Stem Cell Oversight Committee.

## Consent for publication

Not applicable.

## Availability of Data and Material

All relevant EV data was submitted to the EV-TRACK knowledgebase (Van Deun J, et al. *EV-TRACK: transparent reporting and centralizing knowledge in extracellular vesicle research.* Nature methods. 2017;14(3):228-32). To access these parameters use URL: http://evtrack.org/review.php with the EV-TRACK ID (EV230030) and the last name of the first author (Gordillo-Sampedro).

RNA and miRNA sequencing results are available at GEO using accession number GSE229016. The secure token for reviewers is: uxebcoocnjwrrsj

## List of Supplementary Files

Supp Table 1 - miRNA counts table

Supp Table 2 - DESeq2 results for enriched/depleted EV miRNAs

Supp Table 3 - miRNAs with EXOMotifs from García-Martin et al 2022

## Author’s Contributions

Conceptualized and designed experiments (SGS, LA, BL, MM, SAK, FMSDV, AZ, JE), performed and analyzed experiments (SGS, LA, WW), wrote the paper (SGS, LA, AZ, JE). Revised and approved the paper (all authors), supervised and obtained funding (SAK, FMSDV, AZ, JE).

## Acknowledgements

We thank The Centre for Applied Genomics for RNASeq, the SickKids Imaging Facility, Flow Cytometry Facility, and the Nanoscale Biomedical Imaging Facility. We are indebted to C. Smith and J. Rutka (Brain Tumour Research Centre) and thank Sabine Cordes for suggesting EXOmotif searches.

## References

1. Schiweck, J., Eickholt, B. J. & Murk, K. Important Shapeshifter: Mechanisms Allowing Astrocytes to Respond to the Changing Nervous System During Development, Injury and Disease. Front. Cell. Neurosci. 12, (2018).

2. Santello, M., Toni, N. & Volterra, A. Astrocyte function from information processing to cognition and cognitive impairment. Nat. Neurosci. 22, 154–166 (2019).

3. Profaci, C. P., Munji, R. N., Pulido, R. S. & Daneman, R. The blood–brain barrier in health and disease: Important unanswered questions. J. Exp. Med. 217, 1–16 (2020).

4. Houades, V., Koulakoff, A., Ezan, P., Seif, I. & Giaume, C. Gap junction-mediated astrocytic networks in the mouse barrel cortex. J. Neurosci. 28, 5207–5217 (2008).

5. Stogsdill, J. A. et al. Astrocytic neuroligins control astrocyte morphogenesis and synaptogenesis. Nature 551, 192–197 (2017).

6. Patel, M. R. & Weaver, A. M. Astrocyte-derived small extracellular vesicles promote synapse formation via fibulin-2-mediated TGF-β signaling. Cell Rep. 34, 108829 (2021).

7. Ip, J. P. K., Mellios, N. & Sur, M. Rett syndrome: Insights into genetic, molecular and circuit mechanisms. Nat. Rev. Neurosci. 19, 368–382 (2018).

8. Tillotson, R. & Bird, A. The Molecular Basis of MeCP2 Function in the Brain. J. Mol. Biol. 432, 1602–1623 (2020).

9. Albizzati, E. et al. Identification of Region-Specific Cytoskeletal and Molecular Alterations in Astrocytes of Mecp2 Deficient Animals. Front. Neurosci. 16, 1–17 (2022).

10. Sun, J. et al. Mutations in the transcriptional regulator MeCP2 severely impact key cellular and molecular signatures of human astrocytes during maturation. Cell Rep. 42, 111942 (2023).

11. Williams, E. C. et al. Mutant astrocytes differentiated from Rett syndrome patients-specific iPSCs have adverse effects on wildtype neurons. Hum. Mol. Genet. 23, 2968– 2980 (2014).

12. Ehinger, Y. et al. Analysis of Astroglial Secretomic Profile in the Mecp2-Deficient Male Mouse Model of Rett Syndrome. Int. J. Mol. Sci. 22, 4316 (2021).

13. Caldwell, A. L. M. et al. Aberrant astrocyte protein secretion contributes to altered neuronal development in multiple models of neurodevelopmental disorders. Nat. Neurosci. 25, 1163–1178 (2022).

14. Yáñez-Mó, M. et al. Biological properties of extracellular vesicles and their physiological functions. J. Extracell. Vesicles 4, 27066 (2015).

15. Mulcahy, L. A., Pink, R. C. & Carter, D. R. F. Routes and mechanisms of extracellular vesicle uptake. J. Extracell. Vesicles 3, 24641 (2014).

16. Antounians, L. et al. The Regenerative Potential of Amniotic Fluid Stem Cell Extracellular Vesicles: Lessons Learned by Comparing Different Isolation Techniques. Sci. Rep. 9, 1–11 (2019).

17. Théry, C., et al. Minimal information for studies of extracellular vesicles 2018 (MISEV2018): a position statement of the International Society for Extracellular Vesicles and update of the MISEV2014 guidelines. J. Extracell. Vesicles 7, (2018).

18. O’Brien, K., Breyne, K., Ughetto, S., Laurent, L. C. & Breakefield, X. O. RNA delivery by extracellular vesicles in mammalian cells and its applications. Nature Reviews Molecular Cell Biology 21, 585–606 (2020).

19. Pistono, C., Bister, N., Stanová, I. & Malm, T. Glia-Derived Extracellular Vesicles: Role in Central Nervous System Communication in Health and Disease. Front. Cell Dev. Biol. 8, 1–16 (2021).

20. Upadhya, R., Zingg, W., Shetty, S. & Shetty, A. K. Astrocyte-derived extracellular vesicles: Neuroreparative properties and role in the pathogenesis of neurodegenerative disorders. J. Control. Release 323, 225–239 (2020).

21. Sharma, P. et al. Exosomes regulate neurogenesis and circuit assembly. Proc. Natl. Acad. Sci. U. S. A. 116, 16086–16094 (2019).

22. Pan, W., Xu, X., Zhang, M. & Song, X. Human urine-derived stem cell-derived exosomal miR-21-5p promotes neurogenesis to attenuate Rett syndrome via the EPha4/TEK axis. Lab. Investig. 101, 824–836 (2021).

23. Clarke, L. E. & Barres, B. A. Emerging roles of astrocytes in neural circuit development. Nat. Rev. Neurosci. 14, 311–321 (2013).

24. McCready, F. P., Gordillo-Sampedro, S., Pradeepan, K., Martinez-Trujillo, J. & Ellis, J. Multielectrode Arrays for Functional Phenotyping of Neurons from Induced Pluripotent Stem Cell Models of Neurodevelopmental Disorders. Biology (Basel). 11, 316 (2022).

25. Lendemeijer, B., et al. Rapid specification of human pluripotent stem cells to functional astrocytes. bioRxiv (2022). doi:10.1101/2022.08.25.505166

26. Lafourcade, C., Ramírez, J. P., Luarte, A., Fernández, A. & Wyneken, U. MiRNAs in astrocyte-derived exosomes as possible mediators of neuronal plasticity. J. Exp. Neurosci. 2016, 1–9 (2016).

27. Casselli, T. et al. MicroRNA and mRNA transcriptome profiling in primary human astrocytes infected with Borrelia burgdorferi. PLoS One 12, 1–20 (2017).

28. Varcianna, A. et al. Micro-RNAs secreted through astrocyte-derived extracellular vesicles cause neuronal network degeneration in C9orf72 ALS. EBioMedicine 40, 626–635 (2019).

29. Venturini, A. et al. Exosomes from astrocyte processes: Signaling to neurons. Front. Pharmacol. 10, 1–11 (2019).

30. Bartel, D. P. Metazoan MicroRNAs. Cell 173, 20–51 (2018).

31. Nowakowski, T. J. et al. Regulation of cell-type-specific transcriptomes by microRNA networks during human brain development. Nat. Neurosci. 21, 1784–1792 (2018).

32. Squadrito, M. L. et al. Endogenous RNAs Modulate MicroRNA Sorting to Exosomes and Transfer to Acceptor Cells. Cell Rep. 8, 1432–1446 (2014).

33. Garcia-Martin, R. et al. MicroRNA sequence codes for small extracellular vesicle release and cellular retention. Nature 601, 446–451 (2022).

34. Han, K. et al. Human-specific regulation of MeCP2 levels in fetal brains by microRNA miR-483-5p. Genes Dev. 27, 485–490 (2013).

35. Castells, A. A. et al. Unraveling molecular pathways altered in mecp2-related syndromes, in the search for new potential avenues for therapy. Biomedicines 9, 1–22 (2021).

36. Hildebrandt, M. R. et al. Precision Health Resource of Control iPSC Lines for Versatile Multilineage Differentiation. Stem Cell Reports 13, 1126–1141 (2019).

37. Yuan, S. H. et al. Cell-Surface Marker Signatures for the Isolation of Neural Stem Cells, Glia and Neurons Derived from Human Pluripotent Stem Cells. PLoS One 6, e17540 (2011).

38. Cahoy, J. D. et al. A transcriptome database for astrocytes, neurons, and oligodendrocytes: A new resource for understanding brain development and function. J. Neurosci. 28, 264– 278 (2008).

39. Zhang, Z. et al. The Appropriate Marker for Astrocytes: Comparing the Distribution and Expression of Three Astrocytic Markers in Different Mouse Cerebral Regions. Biomed Res. Int. 2019, 1–15 (2019).

40. Semyanov, A., Henneberger, C. & Agarwal, A. Making sense of astrocytic calcium signals — from acquisition to interpretation. Nat. Rev. Neurosci. 21, 551–564 (2020).

41. Antounians, L. et al. Fetal lung underdevelopment is rescued by administration of amniotic fluid stem cell extracellular vesicles in rodents. Sci. Transl. Med. 13, (2021).

42. Lötvall, J. et al. Minimal experimental requirements for definition of extracellular vesicles and their functions: A position statement from the International Society for Extracellular Vesicles. J. Extracell. Vesicles 3, 1–6 (2014).

43. Friedländer, M. R., MacKowiak, S. D., Li, N., Chen, W. & Rajewsky, N. miRDeep2 accurately identifies known and hundreds of novel microRNA genes in seven animal clades. Nucleic Acids Res. 40, 37–52 (2012).

44. Liddelow, S. A. & Barres, B. A. Reactive Astrocytes: Production, Function, and Therapeutic Potential. Immunity 46, 957–967 (2017).

45. Tsuyama, J. et al. MicroRNA-153 Regulates the Acquisition of Gliogenic Competence by Neural Stem Cells. Stem Cell Reports 5, 365–377 (2015).

46. Cho, K. H. T., Xu, B., Blenkiron, C. & Fraser, M. Emerging Roles of miRNAs in Brain Development and Perinatal Brain Injury. Front. Physiol. 10, 227 (2019).

47. Rajman, M. & Schratt, G. MicroRNAs in neural development: From master regulators to fine-tuners. Dev. 144, 2310–2322 (2017).

48. Chu, A. J. & Williams, J. M. Astrocytic MicroRNA in Ageing, Inflammation, and Neurodegenerative Disease. Front. Physiol 12, 826697 (2022).

49. Shenoy, A., Danial, M. & Blelloch, R. H. LetL7 and miRL125 cooperate to prime progenitors for astrogliogenesis. EMBO J. 34, 1180–1194 (2015).

50. Zhang, J. et al. Exosome and exosomal microRNA: Trafficking, sorting, and function. *Genomics*, Proteomics and Bioinformatics 13, 17–24 (2015).

51. Lu, T.-P. et al. miRSystem: An Integrated System for Characterizing Enriched Functions and Pathways of MicroRNA Targets. PLoS One 7, e42390 (2012).

52. Zhang, T. et al. FUS Regulates Activity of MicroRNA-Mediated Gene Silencing. Mol. Cell 69, 787–801.e8 (2018).

53. Grant, C. E. & Bailey, T. L. XSTREME: Comprehensive motif analysis of biological sequence datasets. bioRxiv (2021). doi:10.1101/2021.09.02.458722

54. Kim, Y. J. J., Maizel, A. & Chen, X. Traffic into silence: endomembranes and post-transcriptional RNA silencing. EMBO J. 33, 968–980 (2014).

55. Zhang, Y. et al. Purification and Characterization of Progenitor and Mature Human Astrocytes Reveals Transcriptional and Functional Differences with Mouse. Neuron 89, 37–53 (2016).

56. Khan, M., Hou, S., Azam, S. & Lei, H. Sequence-dependent recruitment of SRSF1 and SRSF7 to intronless lncRNA NKILA promotes nuclear export via the TREX/TAP pathway. Nucleic Acids Res. 49, 6420–6436 (2021).

57. Nasiri-Aghdam, M., Garcia-Garduño, T. & Jave-Suárez, L. CELF Family Proteins in Cancer: Highlights on the RNA-Binding Protein/Noncoding RNA Regulatory Axis. Int. J. Mol. Sci. 22, 11056 (2021).

58. Good, P. J., Chen, Q., Warner, S. J. & Herring, D. C. A family of human RNA-binding proteins related to the Drosophila bruno translational regulator. J. Biol. Chem. 275, 28583– 28592 (2000).

59. Gallo, A., Tandon, M., Alevizos, I. & Illei, G. G. The majority of microRNAs detectable in serum and saliva is concentrated in exosomes. PLoS One 7, 1–5 (2012).

60. Kugeratski, F. G. et al. Quantitative proteomics identifies the core proteome of exosomes with syntenin-1 as the highest abundant protein and a putative universal biomarker. Nat. Cell Biol. 23, 631–641 (2021).

61. Kia, A., McAvoy, K., Krishnamurthy, K., Trotti, D. & Pasinelli, P. Astrocytes expressing ALS-linked mutant FUS induce motor neuron death through release of tumor necrosis factor-alpha. Glia 66, 1016–1033 (2018).

62. Gallo, J.-M. & Spickett, C. The role of CELF proteins in neurological disorders. RNA Biol. 7, 474–479 (2010).

63. Chen, K. et al. miR-125a-3p and miR-483-5p promote adipogenesis via suppressing the RhoA/ROCK1/ERK1/2 pathway in multiple symmetric lipomatosis. Sci. Reports 2015 51 5, 1–15 (2015).

64. Rodrigues, D. C. et al. Buffering of transcription rate by mRNA half-life is a conserved feature of Rett syndrome models. Nat. Commun. 2023 141 14, 1–15 (2023).

65. Cheung, A. Y. L. et al. Isolation of MECP2-null Rett Syndrome patient hiPS cells and isogenic controls through X-chromosome inactivation. Hum. Mol. Genet. 20, 2103–2115 (2011).

