## Supplementary figures and images for "iPSC-derived healthy human astrocytes selectively load miRNAs targeting neuronal genes into extracellular vesicles"

### Supp Figs 1-2

Supplementary Figure 1.

A

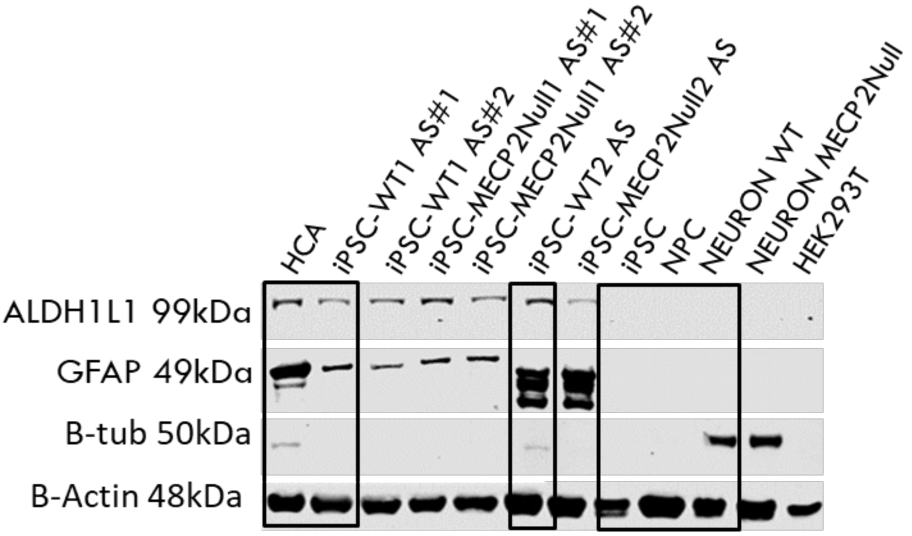


B

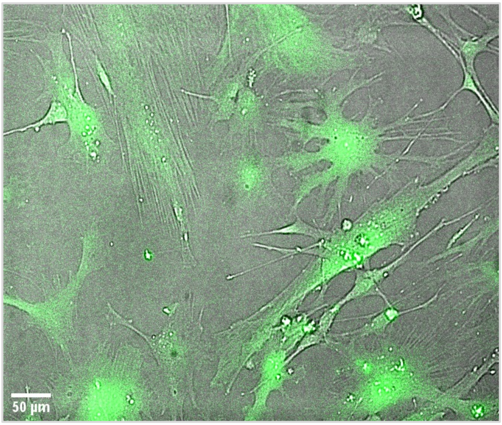


C

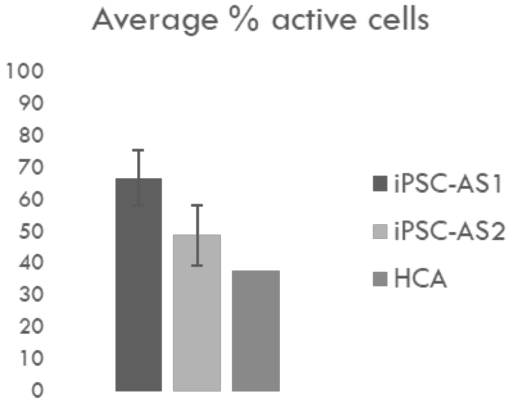

Supplementary Figure 2.

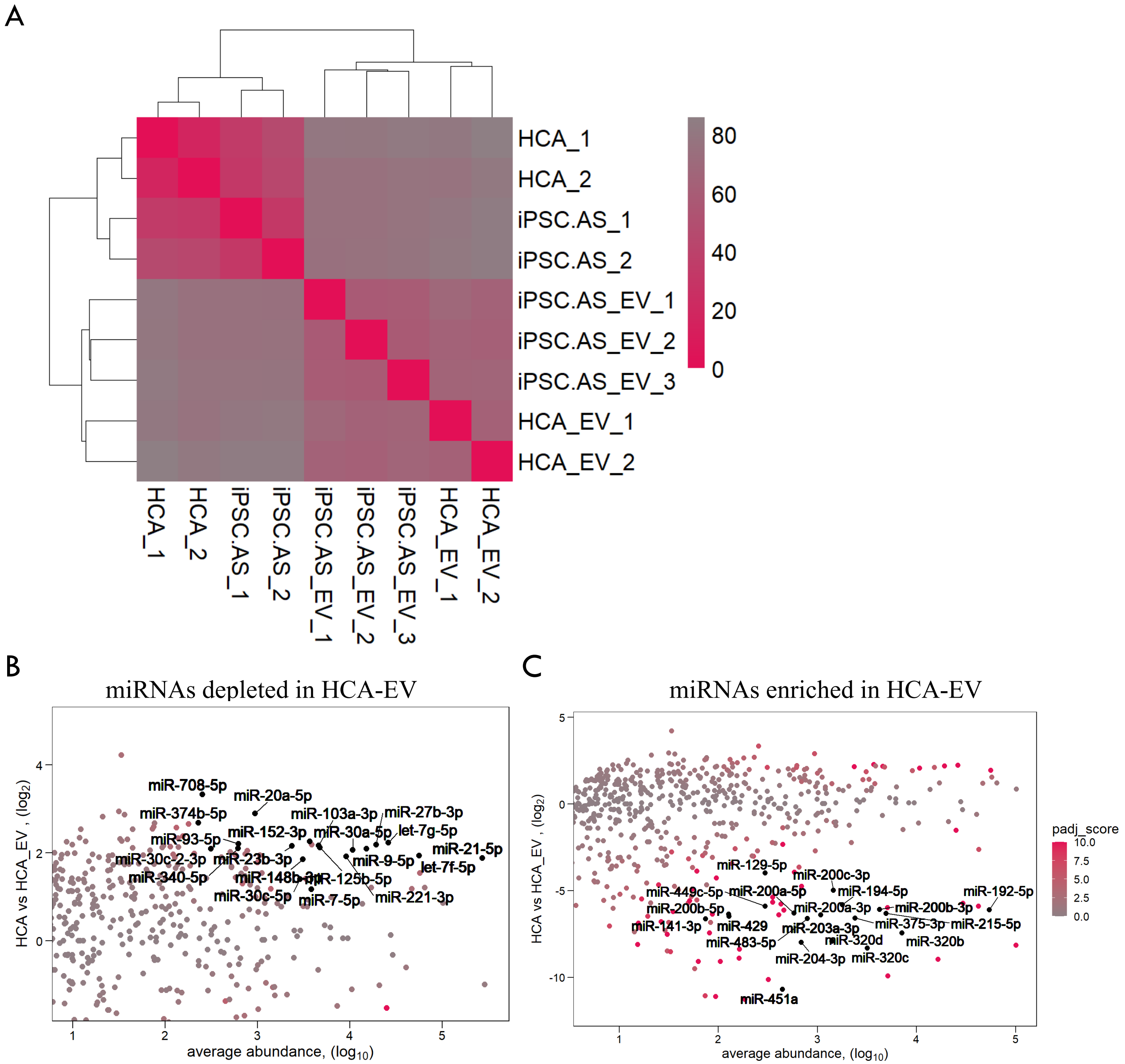
